# Hexokinase and glucokinases are essential for fitness and virulence in the pathogenic yeast *Candida albicans*

**DOI:** 10.1101/448373

**Authors:** Romain Laurian, Karine Dementhon, Bastien Doumèche, Alexandre Soulard, Thierry Noel, Marc Lemaire, Pascale Cotton

## Abstract

Metabolic flexibility promotes infection and commensal colonization by the opportunistic pathogen *Candida albicans.* Yeast cell survival depends upon assimilation of fermentable and non-fermentable locally available carbon sources. Physiologically relevant sugars like glucose and fructose are present at low level in host niches. However, because glucose is the preferred substrate for energy and biosynthesis of structural components, its efficient metabolization is fundamental for the metabolic adaptation of the pathogen. We explored and characterized the *C. albicans* hexose kinase system composed of one hexokinase (CaHxk2) and two glucokinases (CaGlk1 and CaGlk4). Using a set of mutant strains, we found that hexose phosphorylation is mostly assured by CaHxk2, which sustains growth on hexoses. Our data on hexokinase and glucokinase expression point out an absence of cross regulation mechanisms at the transcription level and different regulatory pathways. In the presence of glucose, CaHxk2 migrates in the nucleus and contributes to the glucose repression signaling pathway. In addition, CaHxk2 participates to oxidative, osmotic and cell wall stress responses, while glucokinases are overexpressed under hypoxia. Hexose phosphorylation is a key step necessary for filamentation, that is affected in the hexokinase mutant. Virulence of this mutant is clearly impacted in the *Galleria mellonella* and macrophage models. Filamentation, glucose phosphorylation and stress response defects of the hexokinase mutant prevent host killing by *C. albicans.* By contributing to metabolic flexibility, stress answer response and morphogenesis, hexose kinase enzymes play an essential role in the virulence of *C. albicans.*

**Author summary:** The pathogenic yeast *C. albicans* is both a powerful commensal and pathogen of humans that can infect wide range of organs and body sites. To grow in its host and establish an infection, the pathogen must assimilate carbon from these heterogenous environments. *C. albicans* regulates central carbon metabolism in a niche-specific manner, activating alternatively gluconeogenesis, glyoxylate cycle and the glycolytic metabolism. For yeast and other microorganisms, glucose is the preferred carbon and energy source and its accurate detection and metabolism is essential. However, the glycolytic hexose kinase system has not been investigated yet in *C. albicans.* In this report, we showed that hexokinase and glucokinases contribute to the fitness and virulence of *C. albicans.* We revealed the main metabolic role of the hexokinase CaHxk2 which impacts on growth, glucose signalling, morphological transition and virulence. However, glucokinases contribute to the anoxic response and their implication in regulation processes is suggested.

## Introduction

*C. albicans* is an opportunistic pathogen which exists in a relatively harmless state in the microbial flora of healthy individuals. It is notably present on the mucosal surfaces composing the digestive tract [1, 2]. Perturbations of the normal microbiota, use of medical implants, or predisposing factors like diabetes can trigger *C. albicans* infection. *C. albicans* is the most common cause of fungal nosocomial infections associated with high mortality rates in immunocompromised patients [3-5].

*C. albicans* colonizes diverse host microenvironments such as skin, mucosa, blood, organs, [1]. Among the wide range of virulence traits, survival at 37°C, pH and osmolarity adaptation, secretion of lytic enzymes, alteration of the immune response, morphological changes such as a transition between yeast and hyphae, occur during infection and promote host invasion [6]. Another crucial factor is the metabolic capacity to assimilate host nutrients. The importance of metabolic flexibility to promote systemic infection and commensal colonization has been clearly emphasized during the past years. Genomic tools revealed that rapid transcriptional responses take place to set up a niche-specific carbon metabolism [7-11]. Utilization of alternative non-fermentable sources through the glyoxylate and gluconeogenesis pathways is essential to support *C. albicans* proliferation *in vivo*[12, 13]. However, physiologically relevant hexose sugars like glucose, galactose, and fructose are transiently available at low level in the gastrointestinal tract and only glucose (0.05 to 0.1%) is present in the bloodstream [9, 13, 14]. During colonization of plasma, kidneys and liver by *C. albicans* [14-16], expression of infection-associated genes involved in glycolysis has been reported. Complete glycolytic activation by the two key transcriptional regulators Gal4p and Tye7p is required for full virulence in *Galleria mellonella* and mice [17]. Glucose is the preferred substrate for ATP generation, metabolic precursors synthesis and maintenance of a reductive potential in eukaryotes [18, 19]. Hence, accurate and efficient glucose detection and metabolization pathways constitute a fundamental basis for metabolic adaptation of the pathogen. In *C. albicans*, there are 20 predicted glucose transporters, one of them, CaHgt4, a high affinity sensor of the SRR pathway (Sugar Receptor Repressor), is essential for low glucose level induction of six of the *C. albicans* transporters. In addition, CaHgt4 is also required for filamentation and contributes to virulence in mice [20, 21].

Once detected, the initial step in glucose utilization is its transport and following activation to a sugar phosphate. Most fungi contain at least two active hexose kinases: glucokinase and hexokinase. In *S. cerevisiae*, the hexokinase Hxk2 is the predominant glucose kinase in cells growing in high glucose conditions [22]. Both enzymes can support growth on glucose but hexokinases and glucokinases can also phosphorylate other hexoses like fructose and mannose [23]. The enzymatic equipment for hexose phosphorylation varies among different yeasts, although no physiological explanation for the differences has been found. A search in the genome of *C. albicans* revealed the presence of two hexokinases *(CaHXKl* and *CaHXK2)* and two glucokinase genes *(CaGLKl* and *CaGLK4).* The hexokinase CaHxk1 does not phosphorylate glucose but GlcNAc, an extracellular carbon source present in the mucous membranes, triggering the transition between yeast and hyphal form [24, 25]. However, the hexokinase CaHxk2 and the glucokinases CaGlk1 and CaGlk4 have not been characterized so far and their respective role in *C. albicans* fitness and virulence have not been investigated yet. Moreover, nothing is known on the enzymatic functions of CaHxk2, CaGlk1 and CaGlk4 and their putative dual regulatory role in glucose repression. The preferential use of glucose by yeasts results from glucose-induced transcriptional repression via the CaSnf1, essential AMP-kinase, which phosphorylates the transcription factor CaMig1 [26, 27]. Based on the *S. cerevisiae* model, CaMig1 should form a necessary complex with the hexokinase CaHxk2 to shuttle in the nucleus and generate the glucose repression signal [28], but this particular step has not been described yet in *C. albicans.*

In this study, we evaluated the contribution of CaHxk2, CaGlk1 and CaGlk4 to the phosphorylation of hexoses and to the glucose repression process. Substantial insights in the functional consequences of hexokinase and/or glucokinases deficiency for *C. albicans*growth, various stress responses, morphological transition and virulence are also proposed.

## Results

### Deciphering the hexose kinase activity in *C. albicans*

One hypothetical hexokinase (*CaHXK2*, GenBank XM_712312) and two hypothetical glucokinase genes (*CaGIKl*, GenBank XM_705084 and *CaGLK4*, GenBank XM_707231) were found in the genomic sequence of *C. albicans* (http://www.candidagenome.org/). Analysis of the *C. albicans HXK2, GLK1* and *GLK4* sequences revealed the presence of a classical hexose kinase conserved domain organized in two regions (http://prosite.expasy.org/): a small and a large subdomain. The small subdomain contains the sugar-binding site of typical hexose kinases:-LGFTFSF/YP-[29]. Protein function prediction (http://bioinf.cs.ucl.ac.uk) revealed that CaHxk2, CaGlk1 and CaGlk4 are involved in phosphate-containing compound metabolic processes, transferase activity and ATP-binding. The localization of these conserved domains is provided in S1 Table [30, 31]. Two nuclear localization sequences have been identified in the hexokinase sequence:-PAQKRKGTFT-(8-17) and-QKRGYKTAH-(405-413). These sequences were not found in the glucokinase sequences. Both glucokinase and hexokinase genes are located on chromosome R (Fig. 1A), spaced by 70 Kbp and oriented in opposite directions. The *CaGLK1* and *CaGLK4* sequences share 98.6% and 99.2% identity at the nucleotide and amino acid level, respectively. Alignment of the *CaGLK1* and *CaGLK4* genomic regions, showed that a high level of identity (98%) spanned from 1500 bp before and after the coding sequences. Within these conserved regions, separated by a few hundred of base pairs, both glucokinase genes are framed by two uncharacterized coding sequences. Alignment of these sequences, spanning the 5’ and 3’ regions of both glucokinases, revealed a level of 95.1% and 98.8% identity, respectively. This strongly indicates that the whole conserved region containing the *CaGLK1* and *CaGLK4* genes has been duplicated and conserved.

**Fig.1.**
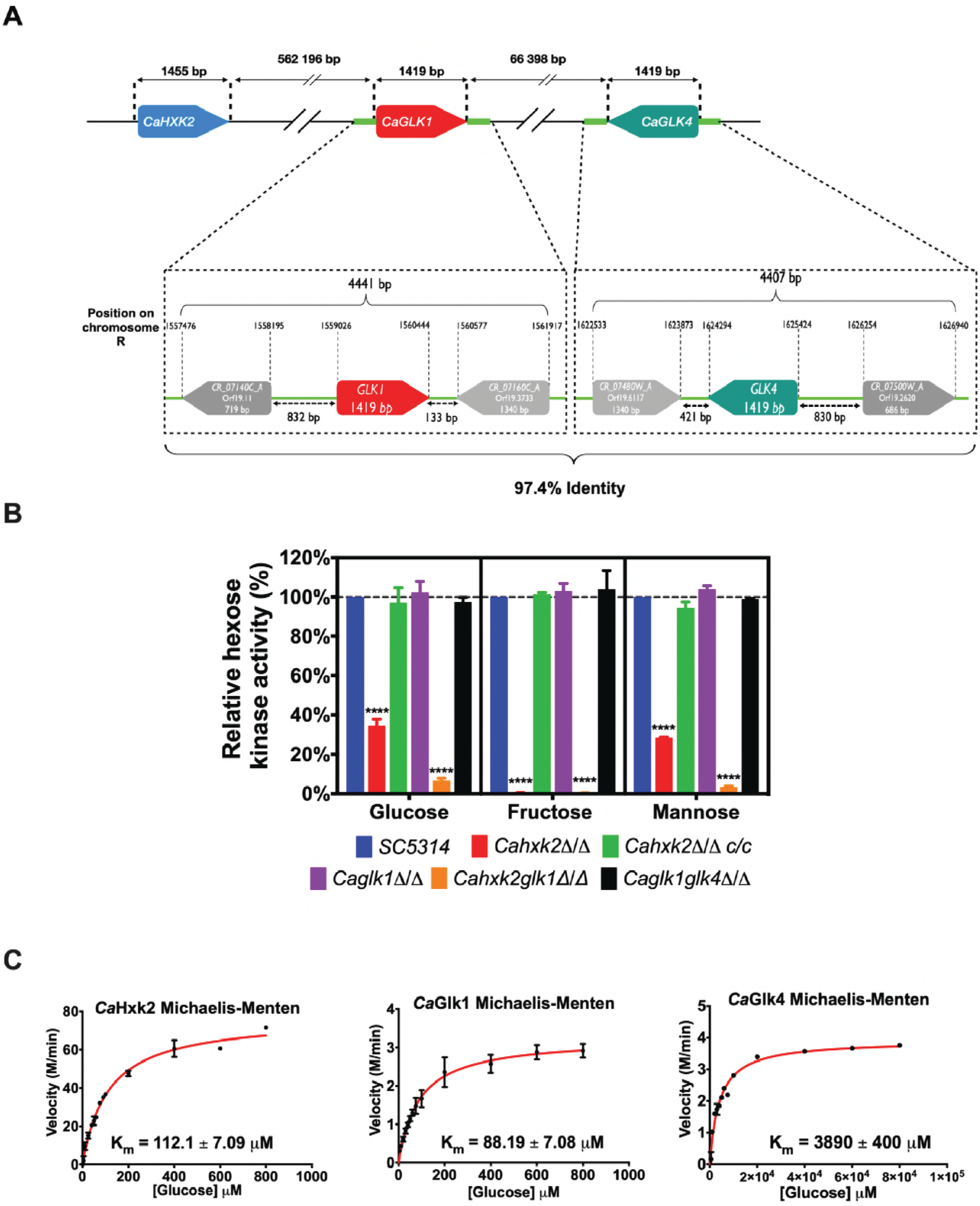
The hexose kinase system in *C. albicans.* (A) The hexose kinase genes are located on *C. albicans* chromosome RA (Ca22chrRA_C_albicans_SC5314:994,376,997,830). Glucokinase genes are oriented in opposite directions and bordered by highly homologous regions (light green). (B) Hexose phosphorylation rates in *C. albicans* wild type and hexose kinase mutant cell extracts. For each strain, the amount of glucose-6-phosphate produced was measured and expressed as a percentage of the wild type strain. Data are presented as a mean (+ standard deviation) of 3 independent experiments performed in triplicates (n = 9); **** P < 0.0001; one-way ANOVA using Dunnett’s method. (C) Kinetic constant of hexose kinases in *C. albicans* in the presence of glucose. The experiment was performed in triplicate. Representative data are presented here.

To identify the function of *CaHXK2, CaGLKl* and *CaGLK4*, a set of gene disruption strains was constructed into the *C. albicans* wild type strain SC5314. Single homozygous null *Cahxk2 [Cahxk2△/△]* and *Caglkl (Caglk1△/△)* mutants, double homozygous null *CaHXK2 CaGLKl (Cahxk2glk1△/△)* and *CaGLKl CaGLK4* (*Caglk1glk4△/△*) mutants were constructed by replacing both wild type alleles using the excisable *CaSAT1* flipper cassette [32]. A *CaHXK2*complemented strain *(Cahxk2△/△ c/c)* was also constructed by reintegrating the wild type coding sequence at the *HXK2* locus, using the same strategy (S1 Appendix).

To investigate the contribution of CaHxk2, CaGlk1 and CaGlk4 to the phosphorylation of hexoses in *C. albicans*, we measured the hexose kinase activity displayed by the wild type strain (SC5314) and the generated mutant strains (Fig 1B). Data obtained with *Cahxk2△/△*cell extracts revealed that glucose kinase and mannose kinase activities decreased by 65 % and 75 %, respectively, while the phosphorylation of fructose was totally abolished. This suggests that other enzymes, like glucokinases, could phosphorylate glucose and mannose, while fructose is phosphorylated by CaHxk2 only. Values obtained with the complemented strain *Cahxk2△/△ c/c*, statistically comparable to the data from the wild type strain, indicated that the lack of fructose phosphorylation was due to the deletion of the gene. Deletion of one or both glucokinase genes *(CaGLK1, CaGLK4)* had no apparent consequence on the level of hexose phosphorylation, suggesting that glucokinase activity could be compensated by the hexokinase activity. Hexose kinase activity measured in the *Cahxk2△/△* strain corresponds to the sum of the activities of CaGlk1 and Caglk4 (33% of the total activity). Glucose kinase activity, measured in the double mutant strain *Cahxk2glk1△/△*, which corresponds to the activity of CaGlk4, was drastically reduced compared to the *Cahxk2△/△*mutant. This activity corresponds to 6% of the total glucose kinase activity. Taken together these results suggest that glucokinases enzymes contribute unevenly and seem to play a minor role in glucose and mannose phosphorylation.

To further investigate the specificity of CaHxk2, CoGIkl and CaGlk4, we determined their apparent Michaelis constant for glucose (Fig 1C). Apparent Km of CoHxk2 was measured in the *Caglk1glk4△/△* strain, while the Km of Glk4 was determined in *Cahxk2glk1△/△* strain. The apparent Km of Glk1 was estimated in the *Cahxk2△/△* by subtracting the effect of CaGlk4. Data revealed that hexokinase 2 and glucokinase 1 have much lower Km values (Km 104.87 ± 7.05 |aM and Km 84.86 ± 6.23 |aM, respectively) than glucokinase 4 (Km 3900 ± 400 |aM), using glucose as a substrate. The low glucose affinity of CaGlk4 could partially explain the poor contribution of this protein to glucose and mannose phosphorylation.

### CaHxk2 mostly sustains growth in the presence of hexoses

Impact of hexokinase and glucokinase gene deletion on growth in the presence of hexoses was evaluated (Fig 2). Delayed growth of the hexokinase mutant *Cahxk2△/△* on glucose and mannose and severely impaired growth on fructose, confirmed the absence of a functional hexokinase. Slow growth on glucose and mannose was consistent with the presence of an additional glucokinase activity. The strong growth defect observed on fructose for this mutant confirmed the fact that fructose is phosphorylated by CaHxk2 only. The residual growth observed on fructose could be due to the metabolism of the alternative carbon sources present in YPG. Growth of the mutant *Cahxk2△/△* was not affected in the presence of glycerol or galactose, substrates that are not phosphorylated by CaHxk2. This indicates that growth defects are linked to an impaired phosphorylation of hexoses. Moreover, growth of the complemented strain was comparable to the wild type. Altogether, these data clearly show that the hexokinase CaHxk2 is necessary for proper growth in *C. albicans.*

**Fig.2.**
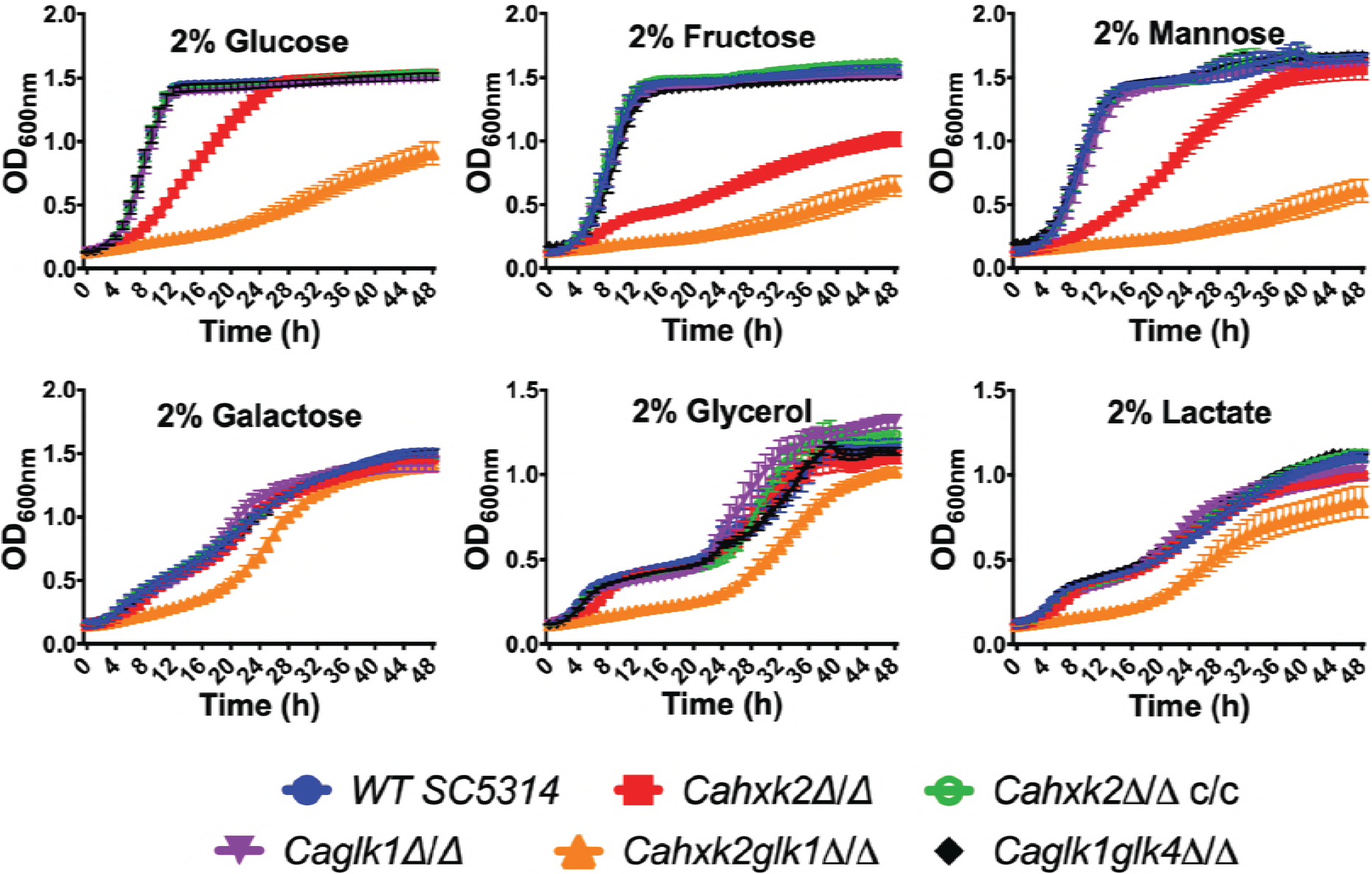
Hexokinase 2 is necessary to sustain *C. albicans* growth. Wild type and mutant strains were grown to the mid-log phase on medium containing 2% of various carbon sources (glucose, fructose, mannose, galactose, glycerol or lactate). A 96-well plate containing appropriate medium was inoculated with each strain at starting OD600nm= 0.2. Cells growth was performed at 30°C and recorded during 48 h using microplates reader (TECAN infinite pro200). Data are presented as a mean (± standard deviation) of 3 independent experiments performed in triplicates (n = 9).

Deletion of *CaGLKl* or both *CaGLKl* and *CaGKLK4* did not affect growth. Growth of the double mutant *Cahxk2glk1△/△* was drastically affected in the presence of glucose, fructose and mannose. Growth failure was also observed, but less pronounced, in the presence of carbon sources that are not phosphorylated by hexokinase or glucokinase (galactose, lactate, glycerol). This strongly suggests that the presence of CoGlk4 alone is not sufficient to sustain growth in the presence of hexoses and that the lack of both CoHxk2 and CoGlk1 could affect general physiological properties, beyond hexose phosphorylation in *C. albicans.*

### Glucokinases and hexokinase do not compensate at the transcriptional level, but exhibit functional redundancy at the protein level

To highlight the respective role of hexokinase and glucokinases, *CaHXK2, CaGLKl* and *CaGLK4* expression was analyzed (Fig 3A). Due to the high level of homology of their coding sequences (98.6 % identity) it has not been possible to amplify *CaGLK1* transcripts alone. Therefore, the transcription level corresponded to the sum of *CaGLK1* and *CaGLK4*transcripts (indicated as *CaGLK1/4).* In the presence of glucose, *CaHXK2* was 3 times more expressed than *CaGLK1/4* (Fig 3A). Transcription of hexokinase and glucokinase genes was strongly induced by glucose (0.1% and 2%). In the presence of glucose, *CaHXK2* was 3 to 5 times more expressed that *CaGLKl/4* (Fig 3A, B). Contrary to glucokinase genes, the level of *CaHXK2* transcripts was dependent on the glucose concentration. Transcription of hexose kinase genes was also strongly induced by mannose and fructose. Surprisingly, the transcription of glucokinase genes was induced by fructose and glycerol which are not substrates for glucokinases (Fig 3B).

**Fig.3.**
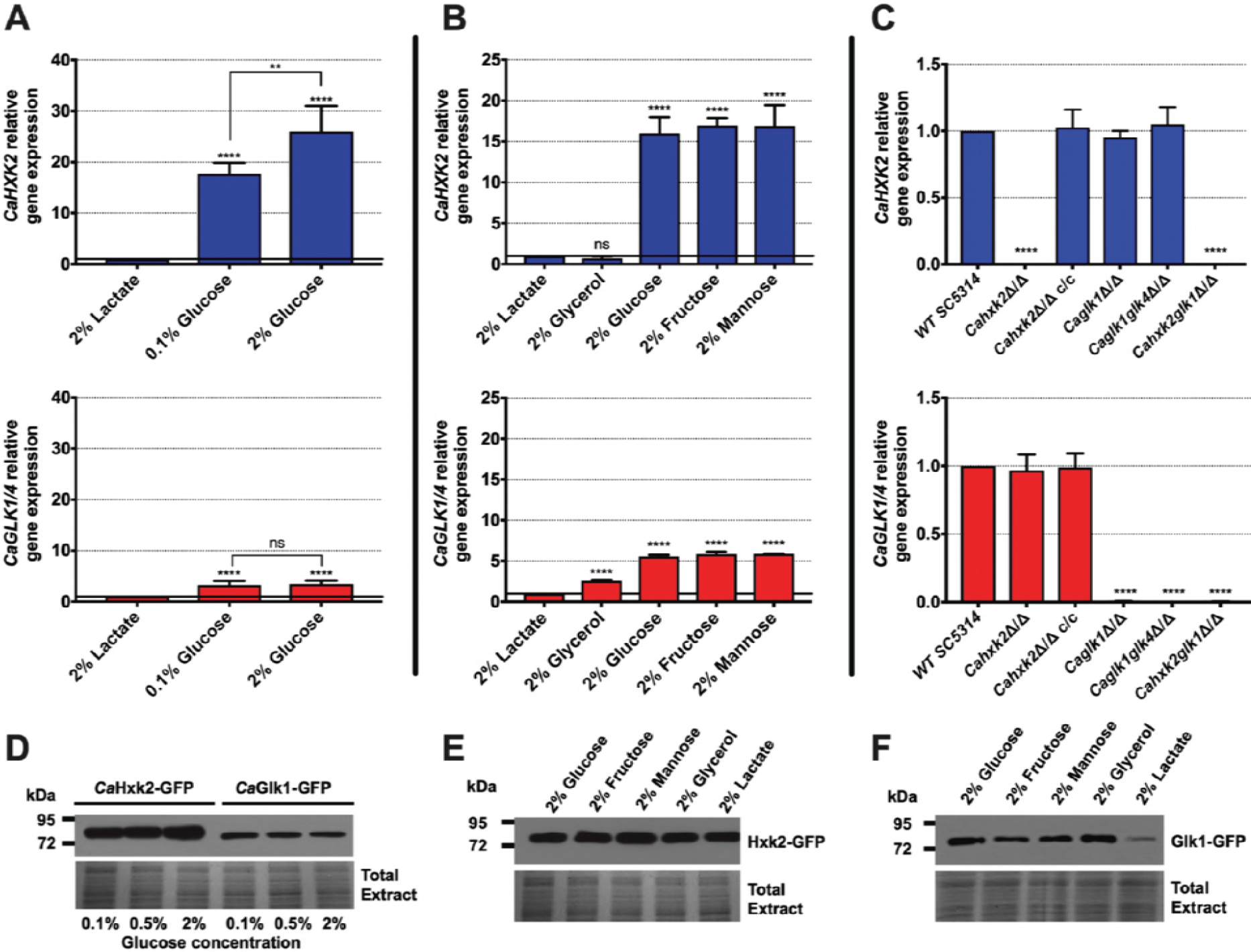
Hexokinase and glucokinases expression. (A) Relative expression of *CaHXK2* and *CaGLK1/4* in wild type strain in presence of 2% lactate, 0.1% and 2% glucose. (B) Relative expression of *CaHXK2* and *CaGLK1/4* in wild type strain in presence of various carbon sources (2% lactate, glycerol, glucose, fructose or mannose). For the panels A and B the results were normalized to the expression of *CaACT1.* The level of *CaHXK2* and *CaGLK1/4*mRNAs was expressed relatively to their abundance in 2% lactate, which was set to 1. (C) Relative expression of *CaHXK2* and *CaGLKl/4* in wild type and hexose kinase mutant strains after growth in 2% YPG. The expression was normalized to the level of the *CaACT1* mRNA internal control. mRNA levels were expressed relatively to their abundance in the wild type strain, which was set to 1. Results represent a mean (+ Standard Deviation) of 3 independent experiments performed in duplicate (n=6); ns, non-significant; ** P = 0.002; **** P < 0.0001. Pvalues were calculated by one-way ANOVA using Tukey’s method. (D) Strains of *C. albicans*expressing C-terminally GFP-tagged CaHxk2 or CaGlk1 were cultivated in presence of 0.1, 0.5 or 2% glucose. Whole cell lysates were analyzed for CaHxk2-GFP and CaGlk1-GFP by Western blotting, using α-GFP antibody. Detection of total proteins by in-gel Coomassie staining was used as a loading control (total extract). Western blots were performed 3 times. Representative data are presented here. (E) and (F) *C. albicans* expressing C-terminally GFP-tagged CaHxk2 or CaGlk1 were cultivated in presence of various carbon sources (2%). Whole cell lysates were analyzed for CaHxk2-GFP and CaGlk1-GFP by Western blotting, using a-GFP antibody. Detection of total proteins by in-gel Coomassie staining was used as a loading control (total extract). Western blots were performed 3 times. Representative data are presented here.

To better elucidate hexose kinase gene regulation, we examined their transcription after growth on 2% glucose in the different mutant strains (Fig 3C). Expression data confirmed an absence of transcripts in the corresponding gene-deleted strains and revealed a complete restoration of the hexokinase transcription level after re-introduction of both wild type alleles. *CaHXK2* expression was not increased in the glucokinase mutants *(Caglkl△/△, Caglk1glk4△/△).* Likewise, *CaGLK1/4* gene expression was not increased in *Cahxk2△/△.* This suggests that unlike what happens in *S. cerevisiae* [33], no compensation mechanisms interfere to regulate glucokinases and hexokinase at the transcriptional level in the absence of one or the other gene. This points out different regulation pathways. However, this compensation could occur at the protein level since the double glucokinase mutant shows no hexose phosphorylation deficiency. Moreover, the fact that the level of *CaGLK1/4* transcripts was unchanged in the absence of the hexokinase *(Cahxk2△/△)* revealed that glucokinases genes are not subjected to glucose repression (Fig 3C). Interestingly, the level of *CaGLK4* expression, detected in *Caglk1△/△* and *Cahxk2glk1△/△* was very low, just above the detection threshold. Considering that the glucokinase gene expression level detected in the mutant *Cahxk2△/△* is the sum of *GLK1* and *GLK4* transcripts, we can again assume that *CaGLKl* and *CaGLK4* are not equally expressed.

To investigate the expression of the enzymes, GFP-tagged CaHxk2 and CaGlk1 were detected in cell extracts by immunoblotting, after growth in the presence of glucose. CaHxk2-GFP and CaGlk1-GFP were detected whatever the glucose concentration. However, CaHxk2-GFP was much more abundant than CaGlk1-GFP (Fig 3D and 3E). This is consistent with the higher transcription level observed for *CaHXK2* but could also reflect a faster turnover for glucokinases. CaHxk2-GFP was equally detected in the presence of various carbon sources that are both inducers of its transcription and substrates of the enzyme, but also and in the presence of glycerol and lactate that do not induce *CaHXK2* transcription (Fig 3B). This could be explained by the long half-life of CaHxk2. In addition to the lowest abundance of CaGlk1-GFP, the main difference between CaHxk2-GFP and CaGlk1-GFP protein content was that CoGIkl was barely detectable in cell extracts after growth on lactate. This may again reflect different regulation processes for CoHxk2 and CoGlk1/4.

### Hexokinase mediates glucose repression but not glucokinases

To highlight the regulatory functions of CaHxk2, we constructed a *HXK2::GFP* strain *(CaHXK2-GFP)* expressing a functional *CaHXK2-GFP* from its own promoter, to examine the localization of CaHxk2 in living cells exposed to glucose (Fig 4A). Upon growth in glucose (2%) the GFP signal was distributed in all the cell (except in the vacuole) with a strong accumulation into a structure that colocalize with the nucleus. This nuclear GFP signal was less important in cells grown in 0.1% glucose and nearly absent in cells grown without glucose (2% lactate) or at very low glucose concentration (0.05%). This indicates that, in *C. albicans*, CaHxk2 is able to shuttle from the cytoplasm to the nucleus in presence of high glucose (0,1% and more). This observation is similar to what observed in *S. cerevisiae* grown in 2% glucose where Hxk2 is known to accumulate into the nucleus where it exerts a transcriptional regulatory function necessary for glucose repression independent of its hexokinase activity [34, 35].

**Fig.4.**
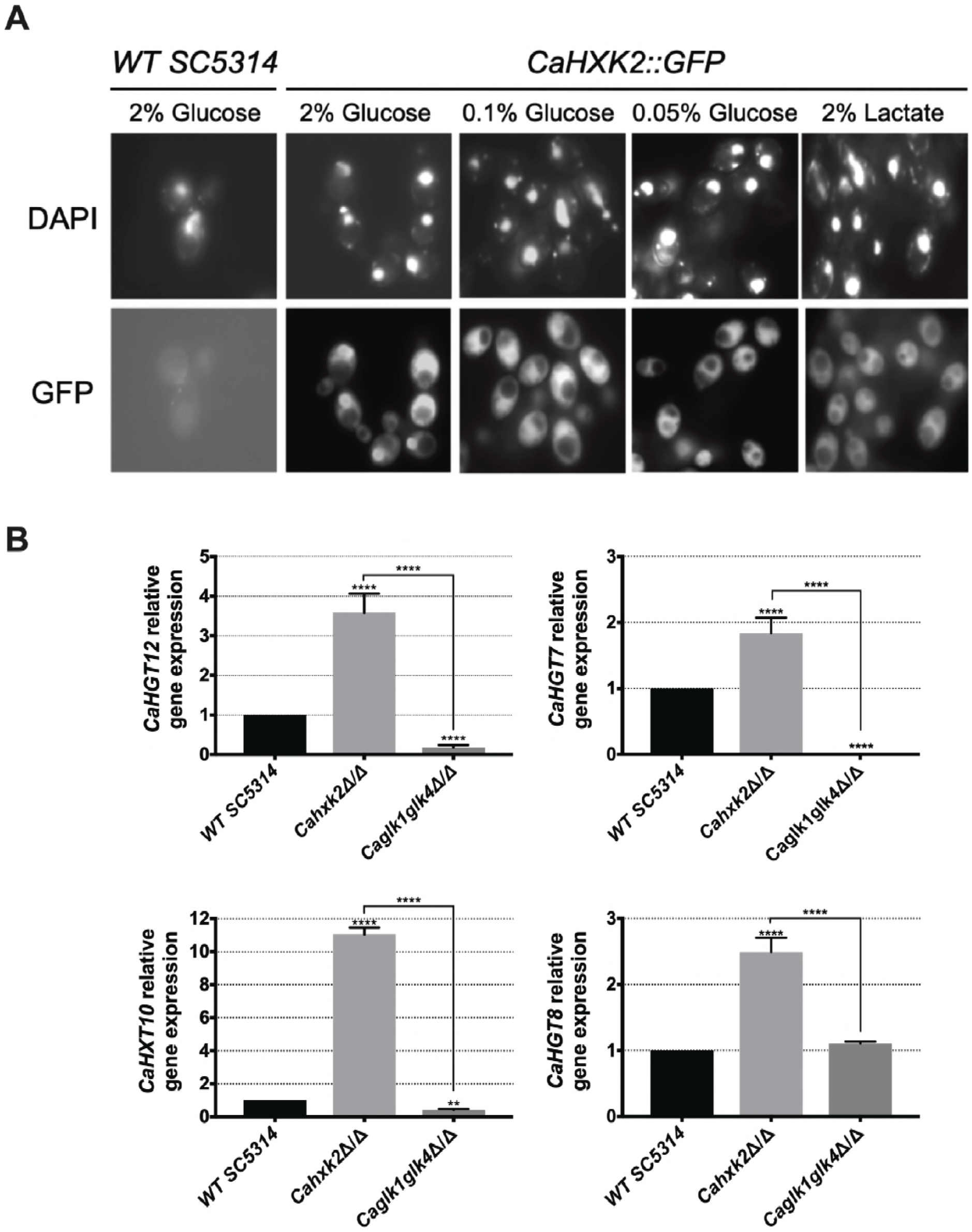
Hexokinase 2, but not glucokinases, participates to glucose repression. (A) Subcellular localization of CaHxk2-GFP was followed using fluorescence microscopy. Direct visualization of CaHxk2-GFP in live cells of *C. albicans* was performed as described in the methods section. Nuclei were identified using DAPI staining. Transformants expressing CaHxk2-GFP were grown on medium containing 2%, 0.1% or 0.05% glucose or 2% lactate as carbon source. GFP and DAPI localization was monitored in live cell cultures using a Zeiss Axioskop 2 Plus fluorescence microscope. Images were taken with a Zeiss AxioCam MR camera using AxioVision software. (B) Expression of glucose permeases, controlled by the Sugar Receptor Repressor pathway *(CaHGT12, CaHXT10* and *CaHGT7)* or not *(CaHGT8)* was measured by qPCR in each strain. Cells were cultivated in the presence of 2% lactate and then transferred for 1 h in 2% glucose before RNA was extracted. Expression levels were normalized to the expression of *CaACT1.* For each permease, the mRNA levels were expressed relatively to their abundance in the wild type strain, which was set at 1. Results represent a mean (+ Standard Deviation) of 3 independent experiments performed in duplicate; (n=6), ** P = 0.002; **** P < 0.0001. Pvalue were calculated by one-way ANOVA using Tukey’s method.

To ascertain the impact of CaHxk2, CaGlk1 and CaGlk4 on glucose repression, we analyzed the expression of high affinity hexose transporter genes that are known to be controlled by the central repressor of the glucose repression pathway, CaMig1, in response to glucose [27, 36] (Fig 4B). These transporter genes *(CaHGT7, CaHGT12, CaHXT10)* are also regulated by another main glucose sensing pathway, the SRR pathway, except *CaHGT8*which is not [20]. Hexose transporter gene expression was drastically enhanced (up to 10 times for *CaHXT10)* in the hexokinase mutant after transfer on 2% glucose medium. On the contrary, *CaHGT7, CaHGT12* and *CaHXT10* expression level was either lowered in the double glucokinase mutant or unaffected in the case of *CaHGT8.* These data suggest that *CaHxk2*but not glucokinases, could have a repressor function on hexose transporter gene expression. Moreover, SRR-dependent transporter genes expression in the glucokinase mutant may suggest an unexpected regulatory role for glucokinase in transporter gene expression.

### Hexose kinase enzymes mediate protection during harmful environmental challenges: glucokinase contributes to the hypoxic response

In *C. albicans* and a number of yeasts, one strategy to counteract oxidative and osmotic stresses is the rapid endogenous synthesis of compatible solutes or, under exposure to cell wall stresses, cell wall biogenesis [37, 38]. These stress responses which are directly or indirectly linked to glucose metabolism, could have been affected in the hexose kinase mutants. For that purpose, wild type and mutant strains were grown in the presence of 2% glucose (YPG) supplemented with 1.2 M KCl (osmotic stress), 5 mM H2O2 (oxidative stress), 0.05% SDS and 5 mM caffeine (cell wall stresses). Data presented Fig. 5A revealed that all stresses had an impact by decreasing growth of the hexokinase mutants *(Cahxk2△/△, Cahxk2glk1△/△).* On the opposite, single and double glucokinase mutant strains were not significantly susceptible to the applied stresses. This suggests that *Cahxk2* is involved in stress responses through its central metabolic position.

**Fig.5.**
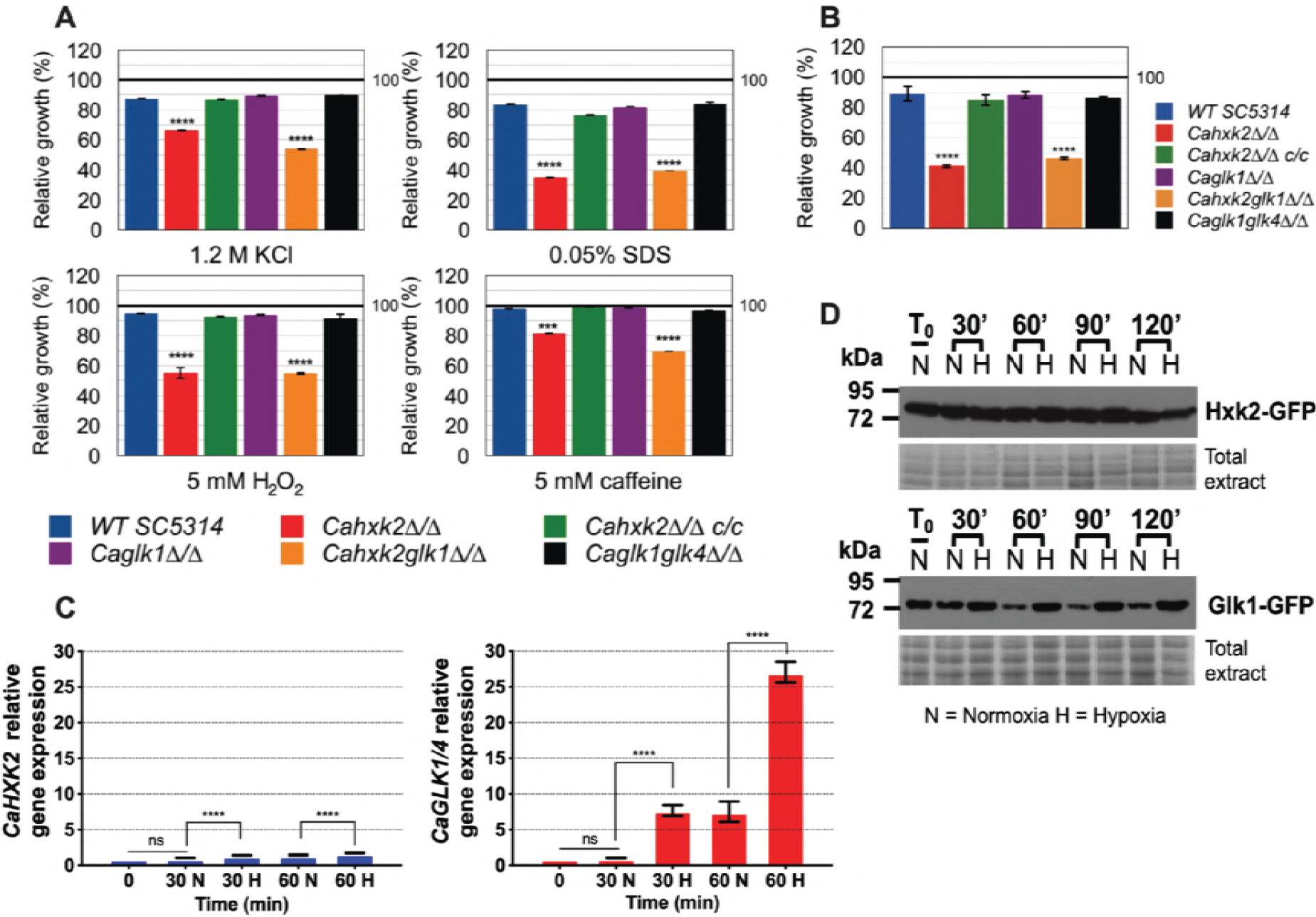
Hexose kinase enzymes mediate protection during harmful environmental challenges. (A) Growth of the wild type and hexose kinase mutant strains in YPG and exposed to various stresses was expressed as a percentage of growth in absence of stress which was set to 100% (black line). (B) Growth of the wild type and hexose kinase mutant strains under hypoxic conditions in 2% glucose YPG was expressed as a percentage of growth in normoxia which was set to 100% (black line). (C) Relative expression of *CaHXK2* or *CaGLK1/4* measured in normoxia (N) or hypoxia (H) during growth in YPG 2% glucose. Transcript level was analyzed by qPCR at 0, 30 and 60 min. Results were normalized to the *CaACTl* transcript level. The level of *CaHXK2* and *CaGLK1/4* mRNAs was expressed relatively to their abundance at time zero, which was set to 1. Histograms represents a mean of 3 independent experiments performed in triplicate (n=9); ns, non-significant; *** P = 0.0002; **** P < 0.0001. Pvalue were calculated by one-way ANOVA using Tukey’s method. (D) Strains of *C. albicans* expressing CaHxk2-GFP or CaGlk1-GFP were grown in 2% glucose YPG to the mid log phase. Cells were transferred into the new medium containing 2% glucose and exposed (n= Normoxia) or not (H= Hypoxia) to oxygen. Following this shift, cells were collected at 0, 20, 60, 90 and 120 min and the detection of CaHxk2-GFP or CaGlk1-GFP was performed by Western Blot using α-GFP antibody. Detection of total proteins by in-gel Coomassie staining was used as a loading control (total extract).

During host infection, *C. albicans* colonizes multiple niches that greatly differ in oxygen content, meaning that it is adapted to hypoxic environments. Growth of the wild type and mutant strains under hypoxic conditions revealed the impact of *CaHXK2* deletion (Fig 5B). Growth of *Cahxk2△/△* and *Cahxk2glk1△/△* was affected by 50% after 24 h, as compared to normoxia while the deletion of one or two glucokinases had minor or no effects. The transcriptional response to hypoxia, elucidated in *C. albicans*, revealed a global upregulation of glycolytic genes [39, 40]. This prompted us to investigate the expression of hexokinase and glucokinase in response to hypoxia (Fig 5C). After one hour of exposure, *CaGLK1/4* transcript level increased by a factor of 25, while *CaHXK2* upregulation was much less detectable. This shows that glucokinases and hexokinase transcription is differently regulated by hypoxic conditions. This was confirmed at the protein level. GFP-tagged hexokinase was equivalently detected in normoxia and hypoxia. In contrast, immunoblot of CaGlk1-GFP revealed a constant amount of protein in response to hypoxia that persisted along the growth, while in normoxia, the amount of CaGlk1-GFP detected clearly decreased (Fig 5D).

### Hexose phosphorylation by CaHxk2 is necessary to fomentation

As glucose is one of the several stimuli that can trigger yeast-to-hypha development in *C. albicans* [41, 42], we checked the ability of hexokinase and glucokinase mutants to undergo a yeast-to-hyphae morphological transition. To evaluate the impact of the hexose phosphorylation step on filamentation, hyphal formation was induced by growth on different media containing known inducing carbon sources, requiring or not hexose kinase enzymes for further metabolization. Spider and serum media, contain respectively mannitol and glucose that depends upon the hexose kinase step to be metabolized. The third medium contains W-acetylglucosamine, that do not require Cahxk2, CaGlk1 or CaGlk4 to be metabolized, but require CaHxk1 [24, 25]. After two days of growth at 37°C on serum and spider media, the wild type, *CaHXK2* complemented strain and glucokinase mutants showed abundant filaments at the periphery of the colony, while the hexokinase mutants *(Cahxk2△/△, Cahxk2glk1△/△)* produced hyphae-deficient colonies (Fig 6A). Morphological changes were also observed at the cell level. Microscopic observations revealed a drastically decreased proportion of filamentous structures for the hexose kinase mutant cells, suggesting that the hexokinase *CaHXK2* is necessary to the yeast-to-hyphae transition. By contrast, filamentation of the *Cahxk2△/△* hexokinase mutant was not affected during growth in the presence of W-acetylglucosamine. All strains behave similarly except the double mutant *Cahxk2glk1△/△*, which appeared hypofilamentous. This suggests that hexose phosphorylation by CaHxk2 could be an essential step for filamentation. Moreover, filamentation defect of the double mutant *Cahxk2glk1△/△* grown on W-acetylglucosamine could be the consequence of severe physiological disturbances.

To eventually highlight a specific role in pathogenic behaviour for *C. albicans* hexose kinases, we compared hexokinase and glucokinase gene expression levels during the early steps of the morphological switch. Our data did not reveal any particular transcriptional response of one gene or another (Fig 6B). Both profiles revealed a two-time increase of transcripts 30 or 60 min after the initiation of the filamentation by serum and a shift at 37°C. However, after one hour of growth, glucokinases expression continues to increase while *CaHXK2* transcription level decreases after 30 min.

**Fig.6.**
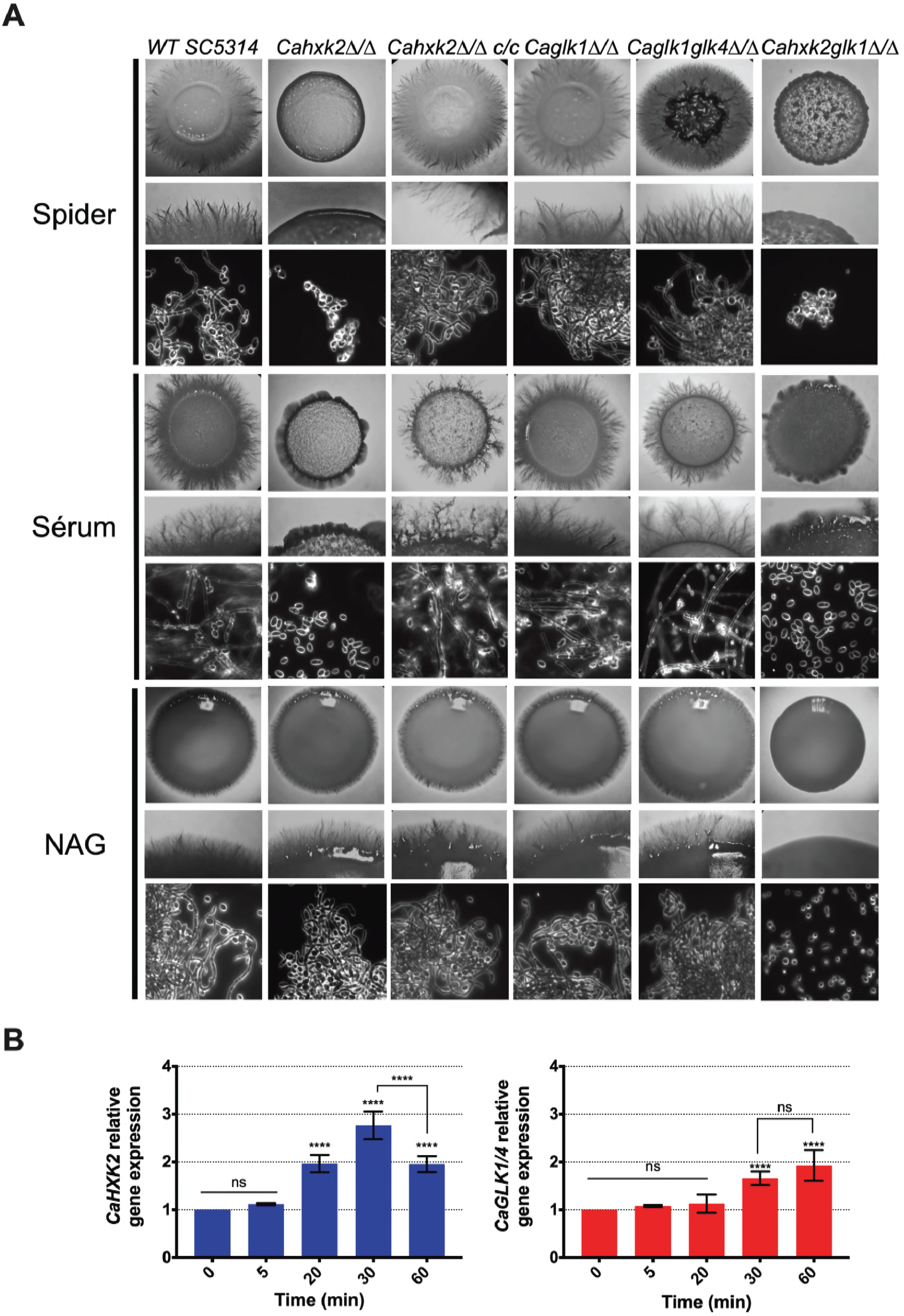
Glycolytic flux is required to sustain hyphal growth of *C. albicans*. (A) *C. albicans* wild type and mutant strains were grown during 3 days at 37°C on spider, serum or N-acetyl-glucosamine medium. For each medium, the upper and middle panels show photographs of macroscopic appearance of the colonies. Photographs present in the lower panel were obtained using Zeiss Axioskop 2 Plus microscope with dark field and show the microscopic aspect. (B) Relative expression of *CaHXK2* and *CaGLK1/4* in the wild type strain during filamentation after transfer from 0.5% YPG to 5% serum liquid medium. Expression level of *CaHXK2* and *CaGLK1/4* was measured by qPCR, at different time points (0, 5, 20, 30 and 60 minutes), and normalized to the level of the *CaACT1* mRNA internal control. mRNAs level was expressed relatively to their abundance at time zero, which was set to 1. Results represent a mean (+ Standard Deviation) of 3 independent experiments performed in duplicate (n=6); ns, non-significant; **** P < 0.0001. Pvalue were calculated by one-way ANOVA using Tukey’s method.

### *Ca*hxk2 mutant is hypovirulent in *Galleria mellonella* and macrophage models

To explore the impact of altering hexokinase and glucokinases on *C. albicans* virulence, we examined first the survival rate of the host model *G. mellonella* following infection with the wild type, mutant and complemented strains (Fig 7A). *G. mellonella* survival data indicated that there was a statistically significant difference between the survival rate of larvae infected by the mutants and the wild type strains, except for the *Caglkl△/△* single mutant. Seven days post infection, 100% of the larvae were killed by the wild type strain while the survival of the larvae infected by *Cahxk2△/△* and *Cahxk2glk1△/△* was still 70% and 85%, respectively. The double glucokinase mutant was also significantly less virulent than the wild type strain, with an intermediary survival rate of 50%. The *Cahxk2△/△ c/c* complemented strain revealed a partially restored virulence.

**Fig.7.**
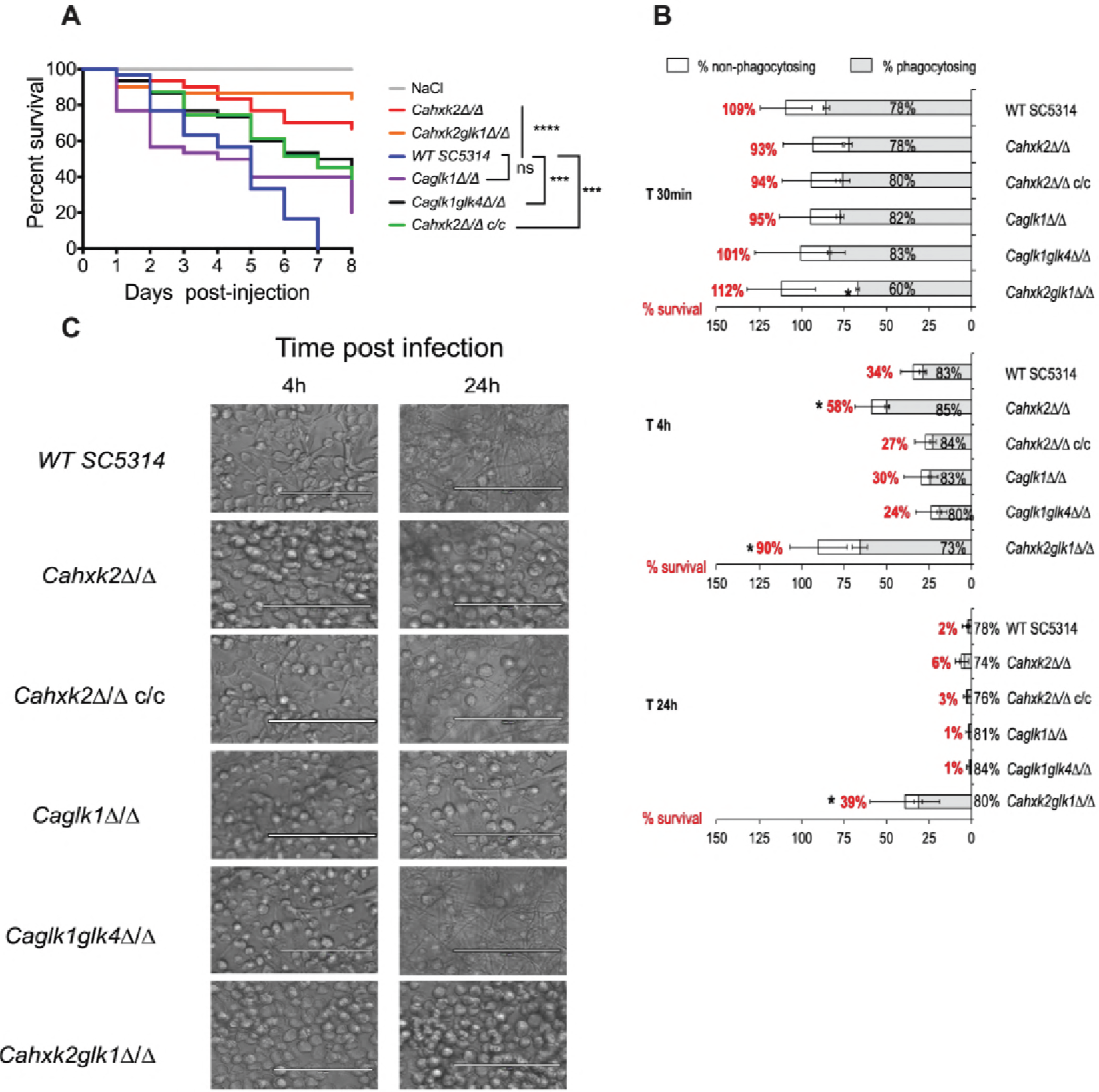
Hexokinase 2 is required for full virulence of *C. albicans.* (A) *Galleria mellonella* model of systemic infection. 2.5 × 10^5^ cells of wild type *(SC5314)*, complemented *(Cahxk2△/△c/c)* or hexose kinase mutant strains were injected into the hemocoel at the last left-pro leg of 30 *Galleria* larvae. Sterile NaCl (0,9%) was injected into control larvae. Survival was monitored for 8 days at 37°C and presented in a Kaplan-Meier plot. Statistical analysis was performed using log rank tests; ns, non-significant; *** P = 0.0002; **** P < 0.0001. *Cahxk2△/△, Cahxk2glk1△/△* and *Caglk1glk4△/△* mutant strains are significantly less virulent compared to the wild type strain (Pvalue ≤ 0.0001). The difference observed between *Caglkl△/△* mutant and the wild type strain is not significant (Pvalue > 0.05). The complemented strain exhibits higher virulence than hexokinase mutants but lesser than the wild type (Pvalue = 0.002). (B) Analysis of mouse macrophage interaction with live *C. albicans* cells in stationary phase at MOI 1:5 (1 macrophage for 5 yeasts) over a 24-hour time course experiment. The horizontal bars represent the macrophage survival, indicated as a percentage on the left side of the bar. The white part of the bars represents the percentage of non-phagocytosing macrophages. The shades tones part represents the percentage of phagocytosing macrophages. (C) Representative pictures of the J774 macrophages after 24 h of infection with wild type and hexose kinase mutant strains in culture flasks at MOI 1:5. The scale bars represent 100

Secondly, we analysed the ability of the mutant to kill macrophages at different interaction times using an *in vitro* model assay (Fig 7B). Hexokinase and glucokinase gene deletions did not modify macrophages association with yeast for any strains, except for the *Cahxk2glk1△/△* double mutant which shows a slightly decreased number of recruited macrophages at the early time of infection (60% compared to approximately 80% for the other strains). This suggests that the absence of hexokinase or glucokinase has no impact on the recognition step. Survival of macrophages was severely enhanced when *Cahxk2△/△* and *Cahxk2glk1△/△* were tested. After 4 hours in the presence of the hexokinase mutant *(Cahxk2△/△)* the number of alive macrophages was nearly twice as high as in the presence of the wild type strain. Moreover, 90% of the macrophages infected by *Cahxk2glk1△/△* were still alive after 4 h, while 34% survived with the wild type strain. After 24 hours, 39% of the macrophages infected with *Cahxk2glk1△/△* survived, compared to only 2% with the wild type strain and 1% for the other strains. This underlines again the very affected virulence capacities of this double mutant. Reintegration of the wild type *CaHXK2* gene restored the killing capacities, suggesting that the virulence defect was linked to the absence of *CaHXK2.*As compared to the wild type strain, interactions performed with glucokinase mutants *(Caglkl△/△, Caglk1glk4△/△)* and macrophages did not reveal significant differences. Because the process of macrophage killing relies on the formation that pierce the phagocytic membrane, the morphogenesis of the strains was analysed at 4 and 24 h post infection (Fig 7C). Our data clearly reveal that *Cahxk2△/△ and Caglk1glk4△/△* did not develop hyphae during macrophage infection. In order to make sure that the growth defect of the hexokinase mutant strains was not the main cause for avirulence, *C. albicans* cells were released from macrophage after 4 hours of phagocytosis by cell lysis and counted (S1 Fig). As compared to the wild type, there was no significant differences in the capacity of the mutant strains to divide inside the macrophage.

Altogether, these data suggest that the virulence defect associated to the deletion of *Cahxk2*could concern the fungal escape phase rather than the recognition and initial phagocytosis steps.

## Discussion

In this study, we sought to assign functions to the hexokinase and glucokinases that could potentially contribute to the fitness and virulence of *C. albicans.* We showed that hexose phosphorylation is mostly assured by CaHxk2, which mainly sustains *in vitro* growth in the presence of hexoses. Hexokinase expression is induced by glucose and higher than glucokinase expression. But proteins are both detectable even in the absence of any phosphorylable hexose. As shown for *S. cerevisiae* glycolytic enzymes, regulation is the result of a complex mixture of gene expression and metabolic effects, in order to optimize simultaneously fluxes, protein and metabolite concentrations [43]. *C. albicans* inhabits niches containing contrasting carbon sources. Metabolic flexibility implies that alternative carbon sources and glucose are assimilated simultaneously [9]. The discrepancy between *C. albicans* transcriptome and proteome has been already clearly highlighted [44]. Upon glucose exposure, CaIcl1 and CaPck1, enzymes involved in the assimilation of alternative carbon sources are not degraded, while their transcripts are subjected to glucose repression. We can assume that a persistent level of CaHxk2, CaGlk1 and CaGlk4 could promote metabolic flexibility and stress response to cope with changing microenvironments reached by the pathogen in the various host niches.

The affected growth profiles of *Cahxk2△/△* and *Cahxk2glk1△/△* indicates the limited ability of CaGlk1 and CaGlk4 to allow glucose utilization in the absence of CaHxk2, while normal growth was observed in the absence of CaGlk1 and/CaGlk4. One possible hypothesis would be a limited glucose uptake caused by the absence of hexokinase. Hence, in *S. cerevisiae* and *K. lactis* glycolytic mutants, glycolysis controls glucose signalling via the SRR pathway. The expression of several glucose-regulated genes, like hexose transporter genes, depends on a functional glycolysis, limiting therefore glucose import [45]. However, this control does not seem to exist in *C. albicans* (Fig. 3C). On the contrary, expression of transporter genes controlled by the SRR pathway *(HGH12, HGT7, HXT10)* was enhanced in the hexokinase mutant *Cahxk2△/△.* Therefore, the poor expression of glucokinase genes, the low intracellular concentration of CaGlk1 and CaGlk4 and their low participation in hexose kinase activity, could mainly explain the growth defect in the absence of hexokinase. Moreover, glycolysis constitutes an interface between metabolism and gene transcription. For instance, glycolysis yields pyruvate which can be oxidized into acetyl-CoA, directly implicated in histone acetylation and gene expression. In stationary yeast cells, increase glucose availability leads to higher levels of acetyl-CoA synthesis, global histone acetylation, accompanied by the induction of a thousand of growth-related genes [46]. The reduced glycolytic flux of the hexokinase mutant could therefore lead to transcription defects and slower growth. In all the tested conditions, the double mutant *Cahxk2glk1△/△* presented an altered phenotype. In this context, CaGlk4 is the only hexose kinase enzyme present. Because of the low affinity of CaGlk4 for glucose and its very low expression, growth of *Cahxk2glk1△/△* is drastically affected in the presence of hexoses. This could be explained by the lack of efficient hexose kinase enzymes. Moreover, hexokinase and glucokinase gene deletions could lead to several drastic intracellular changes. In *S. cerevisiae*, the *hxk2* mutant has a higher H+-ATPase activity and a lower pyruvate decarboxylase activity which coincided with an intracellular accumulation of pyruvate [47]. Absence of glucose repression, could also contribute to redirect carbon flux. In *Klactis*, the identification of hexokinase-dependent proteins related to chromatin remodeling, amino acids and protein metabolism, redox maintenance and stress response reinforces the idea that glucose kinase enzymes exert broader functions than hexose phosphorylation and glucose repression [48].

Our findings reveal that the well-established bifunctional functions of Hxk2 in *S. cerevisiae* [22, 49] also exists in *C. albicans*, while glucokinases do not seem to play a role in glucose repression. We detected CaHxk2-GFP in the nuclei in 0.1% glucose-grown cells (5 mM), which corresponds to the glucose level maintained in the bloodstream and in vaginal secretions [7, 9]. Glucose repression pathway via CaHxk2, could thereby promote metabolic adaptation to favor the fitness of the pathogen, even in glucose-limited host niches. In response to glucose and according to the *S. cerevisiae* model, CaHxk2 should act as a transcriptional carbon catabolite corepressor binding to CaMig1 [50]. *C. albicans* has two orthologs of ScMig1, CaMig1 and CaMig2 but, to our knowledge, no functions have been assigned yet to CaMig2. Transcriptional studies realized on CaMig1 revealed that it regulates a unique set of genes, annotated as carbohydrate uptake and catabolism factors [51]. However, works conducted on CaMig1 revealed that it has no phosphorylation sequence for the kinase CaSnf1, essential for the removal of glucose repression [27, 52]. Deletion of CaMig1 has no effect on the expression of *CaGAL1*, a glucose repressed gene [27] but impacts the transcription of hexose transporter genes [53]. Moreover, CaMig1 has been recently implicated in the resistance to weak organic acids, a novel function [54]. All these evidences show that some of the molecular mechanisms involved in glucose repression in *C. albicans* remain to be elucidated, in particular concerning the direct partners of CaHxk2. Moreover, contrary to *S. cerevisiae* [33] *C. albicans* glucokinases are not subjected to glucose repression. This suggests that glucokinases are not involved into the control of glucose phosphorylation in *C. albicans* and underline their minor role in glucose metabolism.

Contrary to glucokinases, hexokinase gene deletion had an impact on various *in vitro*stress responses. Glucose has been shown to promote stress resistance and to induce some stress genes in *C. albicans* [55]. Our data support this finding, but furthermore indicate a role for glucose phosphorylation in stress resistance. Osmotic and oxidative stresses induce storage of trehalose, glycerol and arabitol [38]. Biosynthesis of such osmolyte sugars and polyols, directly connected to the upper part of the glycolytic pathway, depends on glucose-6-phosphate availability. This was confirmed in *S. cerevisiae* where the analysis of metabolic fluxes in a *Ahxk2* mutant revealed a synthesis of glycerol reduced by a factor of 4.5 [56]. Moreover, the carbon source modulates cell wall architecture and strongly influences the resistance of *C. albicans* to osmotic and cell wall stresses. Glucose and lactate-grown cells display significant differences in cell wall elasticity and ultrastructure [8]. b-glucans are major constituents of *C. albicans* cell wall. Glucan synthases assemble UDP-glucose residues produced from phosphorylated glucose [57]. Thus, any hexose phosphorylation defect would affect the cell wall and render it more sensitive to stress.

Glycolytic gene expression has been associated to the global response of *C. albicans*to hypoxia [58]. Several transcription factors are involved in hypoxia-responsive expression of glycolytic genes. Among them, the key filamentation regulator Efg1 and the transcription factors Tye7 and Gal4 contribute to the early hypoxic response [11, 17, 39, 40]. Contrary to *S. cerevisiae* which ferments sugars even under aerobic conditions, *C. albicans*, a Crabtree-negative yeast, ferments carbohydrates under hypoxic conditions [59]. Our findings specify a clear and drastic effect of hypoxia on glucokinase expression at the mRNA and protein levels. Glucokinase enzymes could be part of the global early hypoxic response, as a spare wheel, to maintain a necessary glycolytic flux during fermentation conditions in oxygen-poor niches. Moreover, this data confirms the fact that hexokinase and glucokinases are not targeted by the same regulatory pathways.

Our results show that the hexokinase mutant retains the filamentation capacity when the carbon source does not require CoHxk2 to be assimilated. Thus, the filamentation-defective phenotype of the hexokinase mutant could be linked to a phosphorylation defect. The absence of one or both glucokinases has no impact on filamentation. This could be related to their low contribution to hexose phosphorylation and, concerning the spider medium, because glucokinases do not phosphorylate fructose. The activities of several major glycolytic enzymes are known to differ in yeast and hyphal forms [60]. We have shown that induction of the filamentation requires upregulation of hexose kinase genes. Morphological switch to filamentous growth requires energy and carbon source, notably to build membranes and cell walls. Thereby, several links between morphogenesis and expression of metabolic genes are established in *C. albicans.* The transcription factor Efg1, part of a Ras-cAMP-PKA signaling network and involved in morphogenesis in *C. albicans*, strongly induces glycolytic and fermentation genes [61]. Moreover, mutants of the CaHgt4/CaRgt1 pathway (SRR pathway) involved in the control of gene expression in the absence of glucose display affected filamentation phenotypes [20, 36]. Thus, nested pathways control simultaneously morphogenesis, glucose signalization and metabolism and by this way CaHxk2, which directly impacts on filamentation through its kinase activity.

The hypovirulence of the hexokinase mutant suggests a central function for CoHxk2. We have shown that the glucose phosphorylation step controls filamentation. Morphological switch is a determining factor of virulence in both *Galleria* and macrophage models. Hyphae are observed in the *G. mellonella* infected tissues [62]. Histological investigations of infected larvae revealed that the SC5314 wild type strain shows a high propensity to filament, leading to gut invasion [63]. Time-lapse microscopy studies have shown a strong correlation between intra-phagocytic hyphal growth and macrophage lysis [64]. Numerous experimental data support a model by which *C. albicans* hyphae enable escape from phagocytes by growing and consequently lysing the cell [65]. CaHxk2 could then be required to sustain hyphal growth within the host cell. Moreover, the impact could be situated at the metabolic level. *C. albicans* hexokinase mutant could degrade trehalose, major hemolymph sugar in *G. mellonella* larvae [66] to recover glucose, but remain unable to phosphorylate it. Recently, Tucey et al., [67] revealed concomitant up-regulation of host and pathogen glycolysis, setting up glucose competition by depleting glucose. *C.* albicans-activated macrophages shift to Warburg metabolism and become dependent on glucose for survival. During macrophage infection, both *C. albicans* free cells and escaped from macrophage could trigger rapid death to the phagocytes by depleting glucose levels. The hexokinase mutant could not compete efficiently for glucose and then turn out to be hypovirulent.

Our data decipher the role of glucose kinase enzymes, not only as a central point of metabolism, but also as actors in regulation, stress response and morphogenesis. Altogether, those different interconnected functions influence the virulence of the yeast. Surprisingly, while the lack of glucokinases did not impact on the phenotype of the mutants, CaGlk1 clearly appeared implicated in the hypoxic response. Moreover, the fact that hexose transporter gene expression level is affected in *Caglk1glk4△/△* suggests that glucokinases could be implicated in regulation processes that remain to be elucidated. Future research might provide further insights in this challenging area.

## Methods

### Strains and media

*C. albicans* strains used in this study are listed in S2 Table. Strains were grown at 30°C or 37°C on YPG medium (1% yeast extract, 2% peptone, 2% glucose). When necessary glucose was added at various concentrations (from 0.01 to 2%) or replaced by other carbon sources like 2% glycerol or 2% lactate. For filamentation assays, *C. albicans* cells were grown for 48 h at 37°C on Spider medium (1% Nutrient Broth, 1% mannitol, 0.2% KH2PO4, 2% agar) or 96 h on YP medium (1% yeast extract, 2% peptone) supplemented with 2.5 mM N-acetyl-glucosamine. Five percent calf serum was also used to induce the morphological switch at 37°C after a 2 to 5 days incubation period. The utilization of different carbon sources and sensitivity to different compounds (5 mM H2O2, 1.2 M KCl, 0.05% SDS, 5 mM caffeine) was monitored in liquid YPG at 30°C by spectrophotometry (Tecan Infinity 200 Pro Serie). Five ml of YPG inoculated with stationary phase cells were cultivated to an optical density at 600 (OD 600) = 0.6. Ten |al of culture were used to inoculate 180 |al of YP medium containing different carbon sources or the required additives distributed in the wells of a plate. Controls lacking carbon source or specific compounds were performed. Plates were sealed with gas-permeable plastic film. OD600 was measured every 30 min during 48 h, with shaking at 380 rpm. Growth data are based on three independent experiments, each of which consisted of assays performed in triplicate. For growth under hypoxic conditions, aerated flasks were inoculated with an overnight culture of *C. albicans*, to OD600 of 0.2. Cells were grown until the beginning of the exponential phase (OD600 ~ 1.8). Hypoxic conditions were created by collecting and transferring cells suspension in hermetic and filled tubes. Different time points following the shift from normoxic to hypoxic growth conditions were considered (30, 60, 90 and 120 min). After appropriate time, cells were collected by centrifugation at 3,000 rpm for 5 min, washed twice with sterile water and rapidly frozen at-80°C. For each time point, three biological replicates were performed. For growth in 96-well plates, anoxic condition were generated by adding 50 |al of mineral oil in each well.

### Construction of mutant strains

Mutant strains used in this study are listed in S2 Table. Mutant strains were constructed using the *SAT1* flipper selection cassette kindly provided by J. Morschhäuser [32]. *CaHXK2*homozygous mutant strain and the complemented strain *Cahxk2△/△c/c* were generated according to methods described by Reub et al., [32]. The *Caglk1△/△* and *Caglk1glk4△/△*homozygous null mutant strains were constructed by one step cloning-free fusion PCR-based strategy. The *Cahxk2glk1△/△* mutant strain was constructed by deleting successively both *CaHXK2* alleles of the *Caglkl△/△* mutant using the *CaHXK2* deletion cassette (S1 Appendix). CaHxk2 and CaGlk1 GFP epitope tagging was performed using a PCR-based strategy using pGFP-NAT1 as a template (kindly provided by S. Bates) [68]. The appropriate mutants were identified by PCR analysis using a combination of primers outside the sites of cassette integration and internal primers.

### Yeast transformation

*C. albicans* transformation was performed using the PEG Lithium technique [69]. After transformation, mixtures were incubated in YPG for 4 h at 30°C and then plated on YPG + nourseothricin 250 |ag/ml (Werner BioAgent, Jena, Germany). Nourseothricin-sensitive cells were obtained according to Reub et al., [32]. Transformants were grown overnight in YPG medium without selective pressure. Cells were plated on YPG containing nourseothricin (25 |ag/ml). Small colonies containing nourseothricin-sensitive cells were selected after 2 days of growth at 30°C. Both alleles were disrupted or complemented in a similar manner after elimination of the *SAT1* flipper cassette. In the case of *in vivo* epitope tagging using pGFP-NAT1 (S3 Table) it was not possible to eliminate the *NAT1* marker, one allele was modified by transformation, only.

### Yeast cell extract and immunoblotting

To prepare proteins extracts, cells were centrifuged and suspended in 500 |al of 0.1 M Tris-HCl buffer supplemented with 10% PMSF (Phenylmethylsulfonyl fluoride). 1.5 ml of glass beads were added and proteins were extracted using FastPrep®-24 (MP Biomedicals). A succession of five grinding (6.5 m/s for 30 sec) was performed. Following this lysis step, cell extracts were centrifuged at 1,500 rpm for 10 min at 4°C. Proteins from the supernatant were quantified using Nanodrop 2000®.

Immunodetection conditions were as described by Rolland *et al.*, [70]. The α-GFP antibody (monoclonal anti-mouse, Roche) and secondary antibody (mouse antibody, HRP conjugated, Bethyl Laboratories) mouse HRP were used at 1/5000^e^ and 1/10000^e^ final concentration respectively.

### Determination of hexose kinase activity

Either glucose, fructose or mannose were used as substrates. The hexokinase II activity was measured spectrophotometrically through NADP+ reduction in a glucose-6-phosphate dehydrogenase-coupled reaction. Each reaction was performed in 1 mL spectrophotometer cuvette at room temperature. The final assay mixture consisted of 100 |al of 25 mM HEPES buffer pH 7.5, 100 |al of 10 mM MgCl2, 100 μl of 1 mM R-NADP, 500 |ag of crude proteins extract, 2 units of Glucose-6-phosphate dehydrogenase and (i) 100 |al of 10 mM D-Glucose for glucose kinase activity, (ii) 2 units of phosphoglucose isomerase and 100 |al of 10 mM D-Fructose for fructokinase activity, (iii) 2 units phosphomannose isomerase, 2 units of phosphoglucose isomerase, and 100 μl of 10 mM D-Mannose for mannokinase activity. Reactions were started with the addition of 100 |al of 5 mM ATP. Absorbance was continuously recorded at 340 nm, for 10-or 15-min. Activities are obtained from the mean of three independent experiments and expressed as a percentage of the activity obtained with wild type crude protein extract. The apparent *Km* of crude extracts of the glucose kinases were determined with a final ATP concentration of 5 mM, a final concentration of glucose ranging from 1 μM to 100 mM. The NADPH apparition at 340 nm was measured using a Tecan Infinite M200 (Salzburg, Austria) microtiter plate reader at 30°C. A single well is composed of 10 μl Glucose-6-P dehydrogenase (0.2 U/mL), 10 |al of 1 mM NADP+, 10 μl of 10 mM MgCl2, 10 μl of HEPES buffer (25 mM, pH 7.6) and 50 |ag of crude extracts. The reaction was initiated by the addition of 10 |aL of ATP (5 mM in potassium HEPES buffer). The activity was determined using a calibration curve of NADH in the range of 0-500 |aM to consider the variability of the optical pathway. The parameters were obtained using Dynafit software and the rapid equilibrium approximation of the Michaelis-Menten equation [71].

### GFP detection by Microscopy

Yeast strain expressing the CoHxk2::GFP and CoGlk1::GFP fusion proteins were grown to exponential phase (OD600 ~ 0.8) in YP containing 0.05, 0.1 or 2% glucose or 2% lactate. Nuclei were stained by addition of DAPI to 10 |ag/ml to the cultures and incubated at 28°C, 180 rpm for 60 min. Cells were washed twice with phosphate buffer saline (PBS) (10 mM Na2HPO4, 1.76 mM KH2PO4, 137 mM NaCl, and 2.7 mM KCl), collected by centrifugation and resuspended in 20 μl of PBS. GFP and DAPI localization were monitored in live cells cultures using a Zeiss Axioskop 2 Plus fluorescence microscope. Images were taken with a Zeiss AxioCam MR camera using AxioVision software and processed using LiveQuartz Images Editor.

### RNA extraction and RT-q-PCR analysis

Total RNA was extracted from cells grown to OD6oo~ 1.5 by the acid phenol method [72]. For reverse transcription-quantitative PCR (RT-qPCR) experiments, 10 |ag of total RNA extract were treated with DNase I (Ambion). Then, ReVertAid H Minus reverse transcriptase (Thermo Scientific), was used as described by the manufacturer, to generate cDNAs. RT-qPCR experiments were performed with the CFX 96 Bio-Rad light cycler using SsoAdvanced Universal SYBR Green Supermix (Bio-Rad). Relative quantification was based on the 2ACT method using *CaACTl* (actin) as calibrator. The amplification reaction conditions were as follows: 95°C for 1 min, 40 cycles of 95°C for 15 s, 60°C for 30 s, and the final step 95°C for 10 s. A melting curve was generated at 95° for 10 s, 65°C for 5 sec with an increment of 0.5 °C until 95°C at the end of each PCR cycle, to verify that a specific product was amplified.

### Infection of *G. mellonella* larvae

For *G. mellonella* infection, overnight cultures of WT (SC5314), mutant or complemented strains of *C. albicans* were grown to stationary phase (OD600 = 5) in 2% YPG medium. Cells were centrifuged and washed three times with 0.9% NaCl. Larvae were infected with 10 μl of suspension (2.5 × 10^5^ cells) injected using a Hamilton syringe, between the third pair of prothoracic legs. Three replicates, each consisting of 10 insects, were carried out with survival rates measured daily for a period of 8 days. Infected larvae were incubated at 37°C in the dark. A control group injected with 10 |al of 0.9% NaCl was included. Death was determined based on the lack of motility and melanisation. Survival curves were plotted and their statistical significance were determined by Kaplan-Meier analysis using the GraphPad Prism 6.0 program. *P* values were estimated using Log rank tests.

### Infection of phagocytes with yeasts

Macrophages from the J774A.1 (ATCC TIB-67) murine cell line were infected as previously described [73] in cRPMI medium (RPMI-1640 without phenol red and supplemented with 10% heat-inactivated fetal bovin serum, 1 mM sodium pyruvate and 2 g/L sodium bicarbonate) at 37°C under 5% CO2. Briefly, 2 × 10^5^ macrophages per well were adhered overnight in 96-well plates, and infected with 1 × 10^6^ Calcofluor White (CFW)-labeled yeast cells in stationary phase in cRPMI medium supplemented with 5 |ag/mL CFW. Interaction was followed over a 24-hour time course experiment. To count yeasts after phagocytosis (S1 Fig), macrophages were infected with *C. albicans* strains as described in Materials and Methods, using 10 macrophages to 1 yeast Multiplicity Of Infection (MOI). After 4 hours of interaction, infected macrophages were collected after trypsin treatment, centrifuged for 10 min at 10000 × g, and lysed in 1 ml of 0.2% ice-cold Triton X-100. Released yeast cells were resuspended in YPD and counted using Kova slides (Kova International). Triplicates were done for each experiment. The results are shown as the average of five independent experiments +-standard errors.

### Flow Cytometry analysis

Flow Cytometry assays were conducted as previously described [73] using a FACSCantoII (Becton Dickinson). Macrophage viability and the ratio of macrophages engaged in phagocytosis were determined after 30 min, 4 h and 24 h of infection with CFW-labeled yeasts. Quintuplets of each condition were done for each experiment. After trypsin treatment, macrophages were labeled with 0.2 μg/mL anti-mouse CD16-APC (a membrane stain) and 0.2 |aM calcein-AM (Sigma) (a marker of active metabolism). The percentage of macrophage viability was calculated using the number of macrophages positive for both fluorescence (anti-CD16-APC and calcein-AM) when infected with yeasts compared to the control uninfected macrophages. Phagocytosing macrophages were quantified as the percentage of the double-stained macrophages also positive for CFW fluorescence. t-test was used to establish statistical significance with a significance level set at P<0.05.

### Statistical analysis

Experiments were performed at least three times independently. All statistical data were calculated with GraphPad Prism 7 software. For comparisons of multiple groups one-way ANOVA method was used. Significance of mean comparison is annotated as follow: ns, not significant; *P=0.033; **P=0.002; ***P=0.0002; ****P<0,0001.

## Acknowledgements

We are grateful to Jade Ravent for technical assistance. We thank J. Morschhäuser and S. Bates for providing the pSFS2A and pGFP-NAT1 plasmids, respectively. The RT-qPCR experiments were performed thanks to the DTAMB (Développement de Techniques et Analyse Moleculaire de la Biodiversite, Universite Lyonl). Romain Laurian was the recipient of a fellowship from the Ministère de la Recherche of France.

## Supporting information

**S1 Table.** Glucose and ATP binding sites and structural domains of yeast hexokinases and glucokinases.

**S2 Table.** *C. albicans* strains used in this study.

**S3 Table.** Plasmids used in this study.

**S4 Table.** Primers used in this study.

**S1 Appendix.** Construction of *CaHXK2* deletion and complementation cassettes.

**S1 Fig**. *C. albicans* wild type and mutant strains divide equally during macrophage phagocytosis.

